# Experts fail to reliably detect AI-generated histological data

**DOI:** 10.1101/2024.01.23.576647

**Authors:** Jan Hartung, Stefanie Reuter, Vera Anna Kulow, Michael Fähling, Cord Spreckelsen, Ralf Mrowka

## Abstract

AI-based methods to generate images have seen unprecedented advances in recent years challenging both image forensic and human perceptual capabilities. Accordingly, they are expected to play an increasingly important role in the fraudulent fabrication of data. This includes images with complicated intrinsic structures like histological tissue samples, which are harder to forge manually. We use stable diffusion, one of the most recent generative algorithms, to create such a set of artificial histological samples and in a large study with over 800 participants, we study the ability of human subjects to discriminate between such artificial and genuine histological images. Although they perform better than naive participants, we find that even experts fail to reliably identify fabricated data. While participant performance depends on the amount of training data used, even low quantities result in convincing images, necessitating methods to detect fabricated data and technical standards such as C2PA to secure data integrity.

## Introduction

Recent years have seen a steady increase in papers retracted by scientific journals, both in absolute terms as well as in proportion to papers published (1). Last year alone saw a record of more than 10,000 papers retracted, with most cases being due to scientific misconduct (1). Actual fraud, the most severe form of scientific misconduct, results in huge follow-up costs: For scientists, funding organizations and companies these take the form of time and resources wasted trying to reproduce fabricated results or the failure of entire projects which were unknowingly based on fraudulent publications (2–6). And, in biomedical fields, these costs can mean inadequate or outright harmful treatments for patients during clinical trials or even clinical practice (5–8).

A common form of fraud in publication, which has become more frequent since years, is the manipulative use of images which are supposed to exemplify or represent experimental results (2, 9, 10). Traditionally, these include inadequate duplication and reuse of images, or manipulations such as the addition, replication, removal, or transformation of objects within an image (2, 10–12). In contrast, modern AI-based methods such as Generative Adversarial Networks (GANs) and Diffusion Models can completely synthesize entire images from scratch (13–15), and are expected to be much harder to detect than previous forms of manipulation (16, 17). This also includes complex images such as histological tissue samples which possess an intricate structure and therefore are harder to manipulate manually.

In the present study, we use the stable diffusion algorithm (14), one of the most recent methods of AI-based image generation, to generate artificial images of mouse kidney histological samples; and in a large online survey with more than 800 participants, we test human ability to distinguish between these artificially generated and genuine histological samples.

By specifically recruiting both, participants with and without prior experience with histological images into our study, we show that prior experience with histological images increases the chance of correct classification, but that even with task-specific training, the overall performance is low. Furthermore, using different sets of artificial images, we demonstrate that human ability to detect such images depends on the amount of data used for training of the stable diffusion algorithm and that very small amounts of training data are already sufficient to generate convincing histological images. Finally, we discuss potential first steps to ameliorate the threat which new, generative methods pose to science integrity.

## Results

In order to test the influence of specific, task-relevant training on the ability to distinguish between genuine and artificial images we exploited the fact that, in contrast to almost all other image categories, to which most people are exposed to in varying degrees in their daily life, there is a sharp distinction between people who either never have seen a histological image (“naive”) and those who have considerable previous experience (“experts”) due to training in a biomedical field. To construct a stimulus set consisting of genuine and artificial histological images of the same structures, we obtained images of periodic acid–Schiff (PAS) stained samples of mouse kidney tissue with a central glomerulus. We then used these to create two sets of artificial histological images using the stable diffusion method in the DreamBooth framework (Fig. 1).One set of images is based on 3 (A3), and one based on 15 (A15) genuine images as training data. In an online survey using the LimeSurvey software, we presented a selection of 8 genuine and 8 artificial images. The images were presented sequentially and for each image participants were asked to classify them as “AI-generated” or “genuine”, with the additional option to skip images. In order to obtain naive and expert populations with similar age, educational status, and socio-economic background, we recruited a total of 1021 undergraduate students from German universities for our study. Out of these, 816 completed the survey and were included in our analysis. 526 participants were classified as experts (186 male, 329 female, 4 divers, 7 not specified; age 23.2 ± 5.1 (mean ± SD)), and 290 were classified as naive (136 male, 137 female, 7 diverse, 10 not specified; age 24.5 ± 5.5 (mean ± SD)).

**Figure 1.**
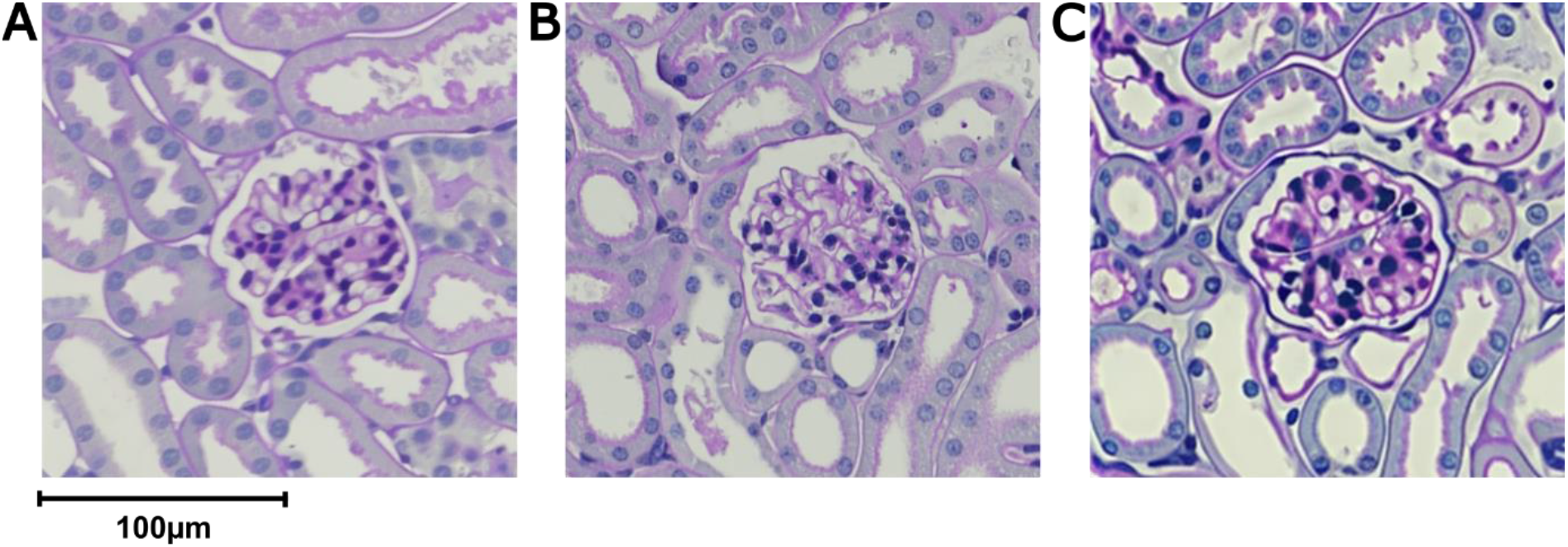
Example images. A) Artificial image created using 15 training images. B) Genuine PAS-stained histological mouse kidney samples images. C) Artificial image created using 3 training images. Images were presented without scalebar during the study.

While we find that, in principle, artificial images can be distinguished from genuine images, this distinction is subject to a high degree of uncertainty (F1 score = 0.64). In addition, we found a clear difference between naive and expert subjects: whereas naive participants only classified 54.5% images correctly (2531/4640 trials, Exact binominal test vs. chance p = 6.2e-10), experts performed significantly better, classifying 69.3% images correctly (5833/8416 trials, Chi^2^ test with Yate’s continuity correction: p < 2.2e-16, Fig. 2A).Overall, both naive and expert participants were able to identify artificial images equally well as genuine ones (Naive: genuine 1245/2320, artificial 1286/2320; expert: genuine 2913/4208, artificial 2920/4208; Chi^2^ test with Yate’s continuity correction: p_naive_ = 0.94, p_expert_ = 0.97). Accordingly, both sensitivity and specificity were higher for expert than for naive participants, whereas within each group sensitivity and specificity were almost identical (Naive: sensitivity 0.55, 95% CI [0.53 0.57], specificity 0.54, 95% CI [0.52 0.56]; expert: sensitivity 0.69, 95% CI [0.68 0.71], specificity 0.69, 95% CI [0.68 0.71]; Fig. 2B).

**Figure 2.**
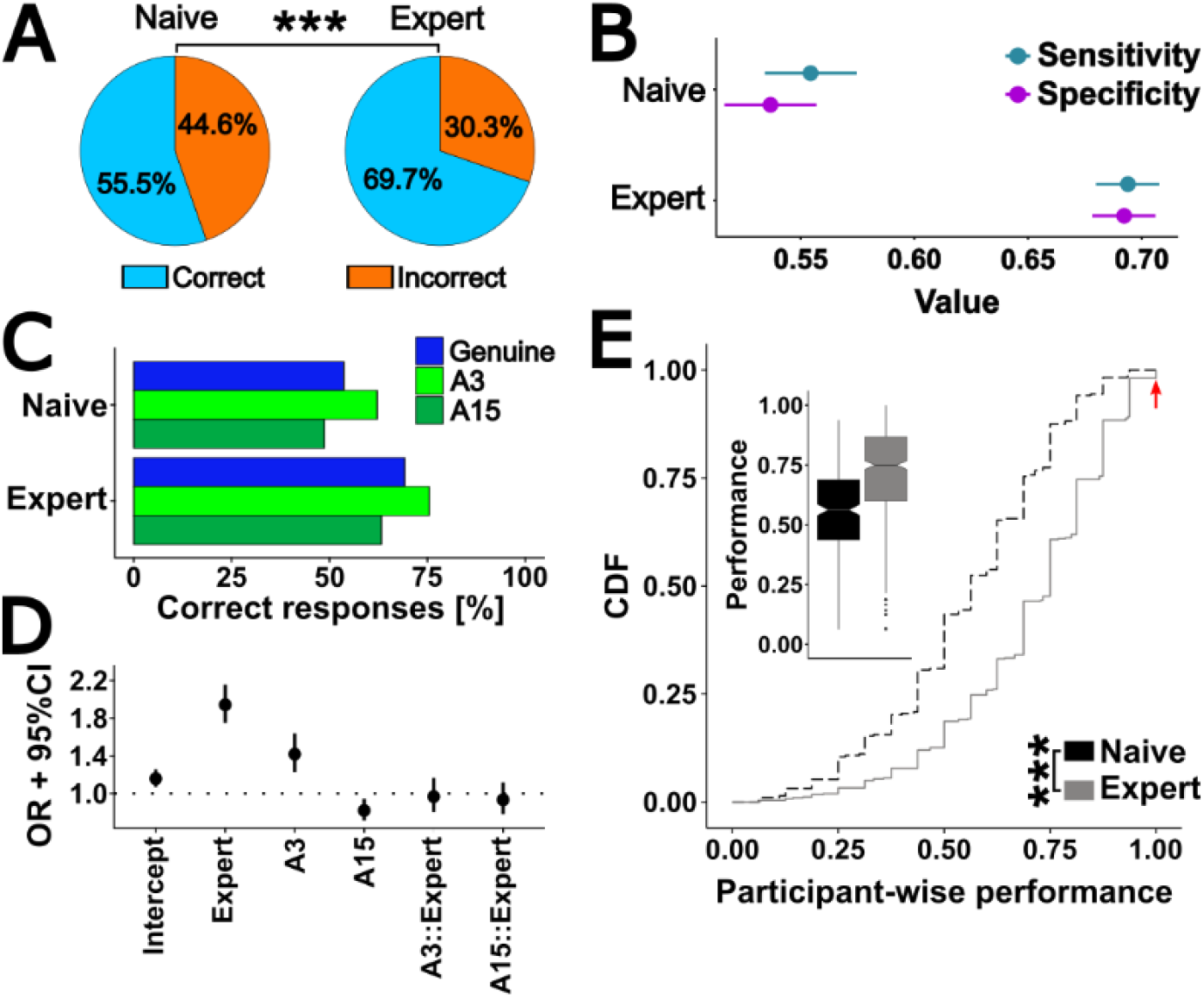
Experts are more successful discriminating genuine from artificial images. A) Overall performance of naive and expert participants (Chi^2^-Test with Yates’ Continuity correction: p < 2.2e-16). B) Sensitivity and specificity for naive and expert participants (point estimates and 95% confidence intervals (CI)). C) Participant performance by image type and prior experience. D) Results for binomial regression on data shown in C, displayed as odds ratio (OR) point estimate with 95% CI. “::” indicates statistical interactions. E) CDF of participant-wise performance by prior experience. Red arrow indicates experts who classified all images correctly. Inset show same data as boxplot. Wilcoxon rank sum test p < 2.2e-16.

However, when we separated artificial images based on the number of images used for training the diffusion model, we found that both, naive and expert participants, classified A3 images more successfully than genuine images (binomial regression, OR 1.42, 95% CI [1.23 1.64], Fig. 2C,D), whereas they classified A15 images less frequently correct than genuine images (OR 0.82, 95% CI [0.71 0.94]). We did not find any significant interactions between participant group and image category (expert::A3 OR 0.97, 95% CI [0.8 1.17], expert::A15 OR 0.93, 95% CI [0.78 1.12]), indicating that the observed effect of image category was not dependent on whether participants had prior experience with histological images.

Importantly, performance was not dependent on stimulus presentation order during the survey (Pearson’s correlation: R2 = 0.058, p = 0.38). In order to test whether the observed difference in performance is present across participants or driven by a few particularly well-performing individuals in the extremes of the distribution, we calculated the individual performance for each participant. Notably, we do find a small group of expert (n =10, 1.9% of all experts) but not naive participants who classified all images correctly. While this performance is outstanding (enrichment ratio against all experts 5.5, Exact binomial test p = 2.0e-5, red arrow in Fig. 2E) even compared to mean expert rather than chance performance (∼70% vs. 50% correct for any image), we observe that experts also perform generally better than naive participants (median participant-wise performance 0.56 vs. 0.75, Wilcoxon rank sum test p < 2.2e-16, Fig. 2E inset). Taken together, our data clearly demonstrate that training enhances the ability to distinguish between genuine and artificial images, but that even experts cannot reliably identify artificial images.

Intriguingly, we found a systematic difference in response times between correct and incorrect classifications, with faster responses for correctly classified images (median 6.99 vs. 7.7s, Fig. 3A). This effect cannot be explained by differences in response times between the two study groups, as experts - who more often classify an image correctly - actually take on average longer for each classification than naive participants (median 6.98 vs. 7.35s; Scheirer-Ray-Hare test: p_correctness_ < 0.001, p_experience_ < 0.001, p_correctness::experience_ = 0.01; Fig. 3B). Instead, we observe the same direction of effect of faster classifications in correct trials in both, naive and expert participants, with astronger difference in experts (median: naive 6.77 vs. 7.38s, experts: 7.07 vs. 8.06s; Wilcoxon rank sum test between correct and incorrect trials with Bonferroni correction for multiple testing: p_naive_ = 0.0015, p_expert_ = 1.2e-16; Fig. 3C). Of note, while almost 100% of response times fall within the first 30s after the trial start, there is an extremely long but thin tail until well over 2000s (Fig. 3A,B). These outliers likely represent cases in which participants started the study, left the browser window and the survey open to do something else, and then finished it at some later time point. To test whether this effect we found could also be observed on a participant-wise basis, we calculated for every participant average response times for correctly and incorrectly classified images, respectively. This paired comparison similarly yielded faster response times for correct classifications, additionally confirming our results on a within-participant level (Median response time 8.33 vs. 9.17s, Wilcoxon signed rank test p = 1.0e-8, Fig. 3D). Taken together, we find that participants classified images faster during correct as opposed to incorrect trials, and that this effect cannot be explained by prior experience with histological images.

**Figure 3.**
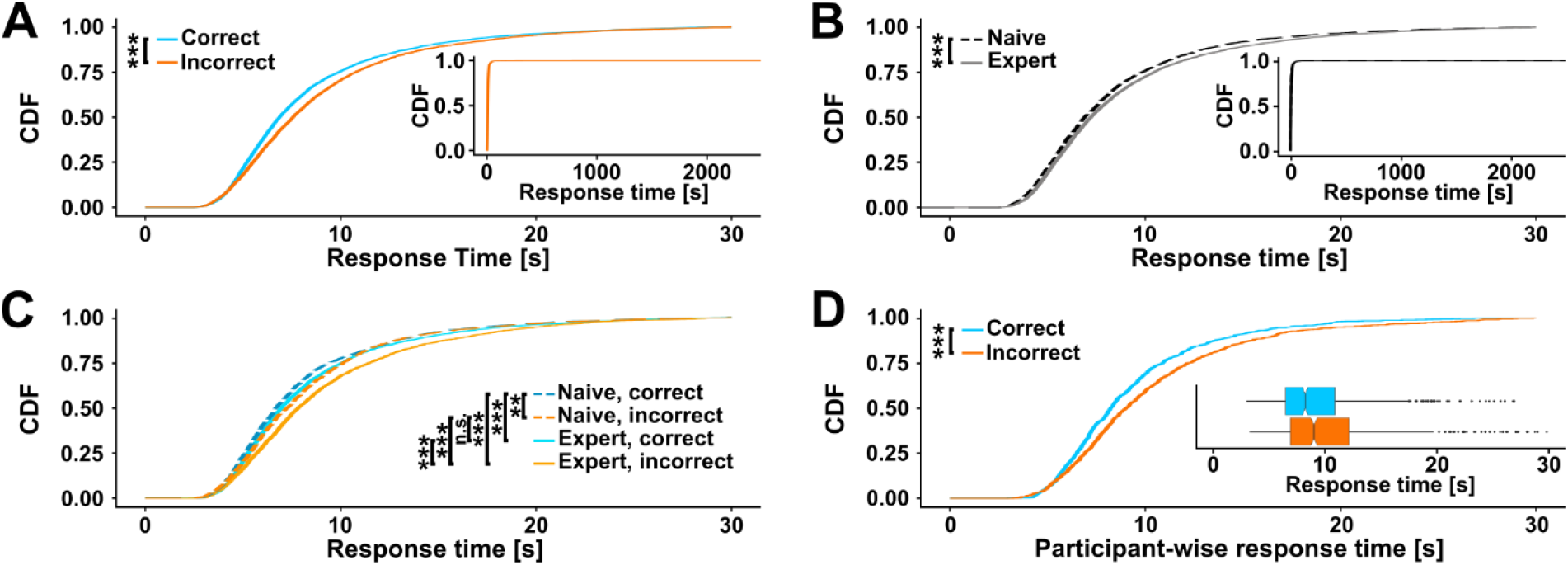
Response times are shorter for correct classifications. A) Empirical cumulative density functions (CDFs) of response times for correctly vs. incorrectly classified images (Scheirer-Ray-Hare test: p_correctness_ <0.001). Inset displays full distributions. B) CDFs of response times for naive vs. expert participants (Scheirer-Ray-Hare test: p_experience_ <0.001). Inset displays full distributions. C) CDFs for group wise comparisons (post-hoc Wilcoxon rank sum test with Bonferroni correction: p_n,c-n,i_ = 0.0015, p_n,c-e,c_ = 2.3e-4, p_n,c-e,i_ = 3.6e-25, p_n,i-e,c_ = 1, p_n,i-e,i_ = 5.0e-10, p_e,c-e,i_ = 1.2e-16). D) CDFs of within-participant response times for correct and incorrect responses (Wilcoxon signed rank test p = 1.0e-8). Inset shows same data as boxplots. Data beyond 30s not shown for display purposes.

## Discussion

In the present study we investigated the ability of human expert and naive participants to distinguish genuine histological samples from artificial ones synthesized using the stable diffusion algorithm (15). We find that naive participants, who have never seen histological images before, are able to classify images better than chance in a statistically significant manner. The actual rate of correct classifications (∼55%), however, was only slightly better than chance (50%). As we did not observe any correlation between stimulus number and performance, it is unlikely that participants learned to distinguish image categories within the study. We assume therefore that the above-chance performance of naive participants results from an overall artificial aspect of synthesized images. While experts performed significantly better than naive participants, overall performance was nevertheless low (70%), demonstrating that distinguishing between real and artificial images is a genuinely hard task.

In addition to these differences in classification performance, we observed that participants were faster when they classified images correctly than when they classified them incorrectly. This effect was not explained by experience with histological images, as it is present within both groups of participants (although more pronounced in experts). In addition, experts, who had a higher rate of correct images, exhibited overall slower response times. As the correctness of a decision is an indicator for its difficulty, faster response times for correct images are in line with canonical models of perceptual decision making, as the drift-diffusion model, in which task difficulty correlates with time allotted to evidence accumulation (18). In contrast, the overall slower response times of experts likely reflect a more active and thus more time-consuming evaluation strategy based on their biomedical training. The performance we observe in our study lies within the range of reported human abilities to discriminate between real and manipulated or synthesized faces (19–21) and natural objects and scenes (22, 23). However, and not surprisingly, in our study the ability to detect artificial images depended strongly on the amount of training data used during image generation, and thus on the resulting image quality. Consequently, as image quality depends both on the precise algorithm employed and the amount of training data, or, in case of manually manipulated images, on the skill and effort of the manipulator, direct, quantitative comparison of the ability to detect forged images between studies should only be done with caution. Furthermore, both sets of artificial images we employed in our study were created based on relatively few training data, preventing a ceiling effect which would make comparisons between groups challenging. The fact that, despite this small training sets in our study and a context which explicitly raised expectations of forged images, suggests that technical hurdles for a forged image to remain undetected in a realistic scenario are low.

This rises practical concerns for the scientific community. It means that readers of scientific articles might easily be fooled, and that neither editors nor reviewers can rely on their personal judgment during peer review. However, though not perfect, viable technical solutions have been proposed, both for scanning scientific publications in particular (2, 24) and for the detection of image manipulation in general (12, 25, 16, 26, 27). Such forensic solutions allow both a higher throughput and consistently outperform humans if applied on the same data sets (19, 23). Where applicable, these methods to detect fraudulent data could be complemented with procedures to authenticate the entire digital workflow including data acquisition, processing, and publication via the implementation of technical standards to secure data provenance such as C2PA (https://c2pa.org). In addition to the introduction of technical measures to promote data integrity, accessibility of the original data associated with any publication is of crucial importance for several reasons: (i) It allows access to high-quality, rather than compressed or otherwise distorted data, which removes obstacles for automated forensic methods (20), (ii) it results in images in their original format and being associated with their original meta data, the combination of which is much harder to fake than an image alone (12, 25), (iii) it is necessary to actually be able to meaningfully enforce the use of standards as C2PA, and (iv) the requirement to submit original data seems to act as a deterrence against fraudulent misconduct *per se* (28).

A very effective time point to perform such an automated scanning is during editorial review of submitted manuscripts (8, 11, 17). Journals can introduce policies that require authors to submit all original data prior to publication, whereas original data may not be recoverable afterwards (9). Similarly, journals can require the application of data provenance standards for scientific data. In addition, methods of image synthesis and manipulation continuously evolve, necessitating a respective development in detection methods (12, 29, 30). Neither reviewers nor readers can be realistically expected to keep up with this development. Similarly, for the particular case of image reuse, journals, but not readers or reviewers can maintain data bases of previously published images for cross-reference (8, 11, 24, 31). Crucially, if original data is checked before, rather than at some distant time point after publication, the enormous follow up costs of fraudulent science which arise for fellow scientist, funding organizations, scientific journals, companies and patients alike can be reduced much more effectively (1, 3–8, 11). And finally, introduction of screening policies by scientific journals has actually been shown to effectively reduce the number of problematic images in accepted manuscripts (11). In conclusion, we demonstrate that it is genuinely hard to spot AI-generated scientific images and that training in the specific field is only somewhat effective at improving the ability to do so; and propose a step towards a solution to ameliorate the threat which new, AI-based methods of image generation pose to science.

## Methods

### Animals

Animal care and experiments were approved by local authorities (Landesamt für Gesundheit und Soziales, Berlin, license G0198/18), and carried out in line with the guidelines of the American Physiological Society. Mice were fed standard rodent chow and provided with food and water *ad libitum*.

### Histology

Male C57BL/6 mice (24-31 g body weight) were sacrificed by cervical dislocation under isoflurane anaesthesia. Kidneys were removed and embedded in paraffin. For Periodic acid-Schiff staining, in paraffin embedded kidneys were sliced in 1.5 µm thin sections and incubated on slides for 16 hours at 60 °C to melt away excessive paraffin. Deparaffinized slices were rehydrated and incubated with 0.5 % periodic acid followed by washing in tab water and incubating with Schiff reagent. After another washing step with tab water, the slices were counterstained with Mayer’s hematoxylin and washed again. The stained slices were dehydrated and mounted with a synthetic mounting medium. Stained kidney slices were recorded using an Eclipse Ti2-A microscope and a DS-Ri2 camera controlled through the NIS-Elements software (Nikon, USA). To obtain large images single images were recorded and stitched afterwards.

### Image generation

Images were generated using stable diffusion 1.5 model checkpoint which has been trained with the DreamBooth (https://github.com/huggingface/diffusers) framework (see code for exact training parameters). The model was trained with either 3 or 15 histological kidney sample images (512×512 pixels, 0,37 µm/pixel). During the survey, the 3 genuine images used to train the model with 3 images, 5 out of the 15 genuine images used to train the model with 15 images, and 4 artificial images from either approach were used.

### Study implementation

The study was conducted online using the LimeSurvey software version 3.25.7+210113 (https://www.limesurvey.org/), hosted at the university Jena and was approved by the local ethics committee (Reference number: 2023-2886-Bef). Participants were recruited from German university undergraduate students by advertisement of the study via student representations, postings in university buildings, or directly in lecture halls. During the survey, 16 images (8 genuine, 8 artificial) were presented, one image at a time. Images were presented without scale bars. For every image, participants were asked to classify the image as “genuine” or “AI-generated”. Participants were familiar with histological images in order to distinguish between naive and expert participants. In addition, we collected data on participant age, gender and response time for every classification. A total of 1021 participants started the study, out of which 816 participants completed the study.

### Data analysis and statistics

All statistical analysis was performed using custom code using RStudio 2023.09.0+463 running R version 4.2.0 (2022-04-22). The versions of all R packages used are documented within the code in the associated repository. Only responses from participants who completed the survey were included in the analysis. If participants did not answer whether they had previously seen histological images (2.8% of all included participants), we assumed they did not know the term and thus treated them as naive in the analysis. If participants skipped single images (1.7% of all included responses), we assumed they were not able to distinguish and treated responses as incorrect trials. For analysis of response times these trials were excluded instead. Crucially, as a sanity check for these assumptions, we additionally performed all analysis excluding both participants who did not indicate whether they had prior experience with histological images, as well as excluding all trials that had been skipped by participants, which yielded the same qualitative and quantitative results. To calculate participant-wise performance, correctly and incorrectly classified images were coded as ‘1’ and ‘0’, respectively, and averaged for every participant. The enrichment ratio was calculated as fold change between the observed number of observations and the number expected under a binomial distribution. To calculate within-participant response time, response times were averaged separately for correct and incorrect trials within each participant, and 10 participants who classified all images correctly were excluded from this comparison. For inferential statistics on frequencies, the exact binomial test was used for comparison to chance levels, and the Chi^2^ test with Yate’s continuity correction was used to test for statistical difference between groups. To test the statistical contribution of prior experience and image category on classification performance, we used a general linear model assuming a binomial distribution and calculated the odds ratio as OR = e^Est^ where ‘Est’ denotes the model point estimates. For inferential statistics on continuous data, data was tested for normal distribution using the Anderson-Darling test. All continuous data sets were non-normally distributed; thus, the Wilcoxon rank sum test was used for non-paired, and the Wilcoxon signed rank test for paired data. To test group effects on response times, we used Scheirer-Ray-Hare test was used as non-parametric alternative to a multi-way ANOVA, and post-hoc pair-wise comparisons were made using the Wilcoxon rank sum test with Bonferroni’s correction for multiple testing. For calculation of sensitivity and specificity, we counted genuine images as negative and artificial images as positive ground truth, and confidence intervals (CI) were calculated using the Wilson score interval method. Statistical levels are indicated in figures as: ‘n.s.’: p ≥ 0.05, ‘^*^’: p < 0.05, ‘^**^’: p < 0.01, ‘^***^’: p < 0.001. ‘::’ indicates statistical interaction between factors. Boxplots indicate median, 25^th^-75^th^ percentile (box) and last data point within 1.5^*^IQR (inter-quartile range) below 25^th^ or above 75^th^ percentile (whiskers). Outliers beyond whisker range are indicated as single data points and were always included in statistical analyses. Boxplot notches indicate the 95% CI of the median and are calculated as median ± 1.58^*^IQR/sqrt(n), where ‘IQR’ denotes the inter-quartile range, ‘sqrt()’ the square root and ‘n’ the number of data points.

## Data availability

Original data collected in this study and images used for the survey will be publicly available on GitHub and zenodo as of date of publication.

## Code availability

All code used in this study will be publicly available on GitHub and zenodo as of date of publication.

## Acknowledgements

We would like to thank all members of the Mrowka lab for discussion of the project. This study was supported by the Deutsche Forschungsgemeinschaft (DFG, German Research Foundation) – Project ID 394046635 – SFB 1365, the Free State of Thuringia under the number 2018 IZN 0002 (Thimedop) and the European Union within the framework of the European Regional Development Fund (EFRE).

## Author contributions

Conceptualization: RM

Methodology (artificial image synthesis, survey design): RM

Methodology (animals, histology): VAK, MF

Formal analysis: JH, CS

Visualization: JH

Investigation (participant recruitment): JH, SR

Writing: JH, RM

Supervision: RM

## Materials & Correspondence

Inquiries regarding image generation or data acquisition should be directed to RM. Inquiries regarding data analysis should be directed to JH.

